# Heterogeneous Slowdown of Dynamics in the Condensate of an Intrinsically Disordered Protein

**DOI:** 10.1101/2024.07.15.603508

**Authors:** Saumyak Mukherjee, Lars V. Schäfer

## Abstract

The high concentration of proteins and other biological macromolecules inside biomolecular condensates leads to dense and confined environments, which can affect the dynamic ensembles and the time-scales of the conformational transitions. Here we use atomistic molecular dynamics (MD) simulations of the intrinsically disordered low complexity domain (LCD) of the human fused in sarcoma (FUS) RNA-binding protein to study how self-crowding inside a condensate affects the dynamic motions of the protein. We found a heterogeneous retardation of the protein dynamics in the condensate with respect to the dilute phase, with large-amplitude motions being strongly slowed by up to two orders of magnitude, whereas small-scale motions, such as local backbone fluctuations and side-chain rotations, are less affected. The results support the notion of a liquid-like character of the condensates and show that different protein motions respond differently to the environment.

---

The discovery of membraneless organelles (MLOs) has changed cellular biophysics,^1^ challenging the traditional view that cellular compartmentalization is exclusively mediated by lipid-bound organelles and vesicles. MLOs are high-density liquid droplets formed through the process of liquid-liquid phase separation (LLPS)^2–6^ of proteins, particularly intrinsically disordered proteins (IDPs),^7,8^ and RNA.^9^ They play critical roles in orchestrating various cellular processes.^3,10–13^ For example, the biochemical reactions occurring in MLO compartments are crucial for transcription, stress response, synaptic activity, RNA splicing, receptor-mediated signalling, and mitosis.^14–22^

Due to their ability to concentrate biomolecules, MLOs are described as biomolecular condensates or biocondensates.^23^ The concentrations of proteins and/or nucleic acids inside the condensates are typically higher than 100 mg mL*^−^*^1^.^24–27^ This leads to increased viscosity inside the droplets, which can be 3 to 6 orders of magnitude higher than in dilute solutions. ^28^ Consequently, the diffusion of biomolecules in condensates is strongly retarded. ^24,25,29–33^ For example, an up to 500-fold decrease of translational diffusion was reported for condensates of the intrinsically disordered low-complexity domain (LCD) of the human fused in sarcoma (FUS) RNA-binding protein.^25^

To fulfil their biological functions, biocondensates need to ensure a high local concentration of certain biomolecules. At the same time, these biomolecules need to retain their dynamic nature as much as possible, and such liquid-like character is also crucial for the exchange of molecules between condensates and their surrounding. The dynamic nature of condensates partly originates from the high degree of conformational flexibility of the constituent IDPs, fostered also by the relatively high water content of the droplets.^25,34,35^ Driven by their plasticity and the lack of a membrane the condensates can rapidly form, reshape and adapt their properties, and hence efficiently interact and exchange molecules with their surroundings.^36^ The exact link between the dynamic nature of condensates and their biological functions can sometimes be difficult to determine, though, and it may strongly depend on the particular system under investigation.^13,37^ In a striking recent example, Fischer et al. showed that transcription factor condensates with more liquid-like character lead to higher gene expression levels than stiffer condensates.^38^

The dynamic and broad conformational ensembles of IDPs, as dictated by their amino acid sequences, lead to dynamic multivalent interactions that play a key role in driving LLPS and stabilizing biocondensates. ^5,39^ In general, the flat free-energy landscapes of IDPs imply lack of a stable persistent conformation, but a highly dynamic conformational ensemble that is characterized by rapid transitions between a multitude of metastable minima. ^40,41^ This plasticity of IDPs^42^ fosters dynamic interactions in the condensates, which includes mechanisms such as folding-upon-binding, formation of highly dynamic biomolecular complexes, conformational selection, and fly casting.^43–46^ Some of these mechanisms involve the transient formation of (local) structure, e.g., via the restriction of certain dynamic modes.^28^ The increased viscosity in highly concentrated condensates is expected to further restrict the conformational dynamics of the proteins, although the droplets retain a liquid-like character and the constituent proteins can still exhibit rapid chain dynamics.^47–49^

Zheng et al. used MD simulations to investigate the multivalent interactions that govern LLPS of FUS-LCD.^47^ They showed that ion partitioning between dense and dilute phases is largely driven by the charge distribution along the protein chain, and that no strong enrichment or depletion of ions is found in condensates due to the small net charge of FUS-LCD.^47^ Furthermore, the MD simulations of Zheng et al. also showed that, despite the high concentration inside the droplets, the protein molecules remain mobile, and that the dynamic self-association of the proteins is driven by a combination of nonspecific interactions as well as hydrogen bonds, salt bridges, and π–π and cation–π interactions.^47^

In a combined experimental/computational study, Galvanetto et al. ^48^ used fluorescence spectroscopy and MD simulations to investigate the ProTα–H1 system. ProTα has ultrahigh affinity towards H1 due to strong electrostatic interactions of the two highly charged proteins (the net charges of ProTα and H1 are -44 and +53, respectively, which strongly differ from the small net charge of FUS-LCD of -2). A main finding of that study was that nanosecond time-scale dynamics are retained in the condensate despite a ca. 300-fold higher viscosity than in the dilute phase, a conclusion that agrees with the results reported by Zheng et al. for FUS-LCD. Interestingly, a recent NMR relaxation study of the measles virus N*_T AIL_* protein, an IDP that undergoes LLPS, showed that while sampling of the protein backbones is not strongly affected in the dense phase, librational, backbone torsional, and segmental (or chain-like) dynamics are considerably slower, ^49^ a finding that needs to be reconciled with the results of Galvanetto et al. Along the same lines, a time-resolved fluorescence spectroscopy study found that α-synuclein has reduced chain flexibility under LLPS conditions compared to dilute solution.^50^ However, no specific dynamic protein modes were identified, and the effects on backbone and side-chain motions were not separately addressed. All in all, we conclude that further studies are needed to characterize and understand the dynamic behavior of biomolecules in condensates in more detail.

Here we approach biomolecular condensates from a fundamental biophysical chemistry perspective and investigate on the effect of the dense environment provided by a biomolecular condensate on the protein dynamics. Single-component condensates formed by the low-complexity domain (LCD) of human FUS are used as a model system to investigate which dynamic modes of the protein are affected how, and to what extent, in the dense condensate environment as compared to dilute conditions. FUS is physiologically important for RNA shearing and transport, DNA repair, micro-RNA processing, gene transcription and regulation.^51–53^ The LCD region is primarily responsible for driving LLPS.^24,25,27,54,55^ It encompasses the N-terminal 163 residues of the protein and is rich in glutamine and serine residues, tyrosines, and also glycines, which are known to be important for condensate formation via LLPS.^56^

All-atom MD simulations were used to study the dynamics of the protein backbone and side-chains of FUS-LCD in condensates and compare them to dilute conditions (snapshots of the systems are shown in Figure S1). To understand the condensate effects on the dynamics of the protein chains, the backbone and side-chain motions were investigated separately. First, the overall translational dynamics of the FUS-LCD chains were characterized by calculating the mean squared displacements (MSDs) of the protein atoms. From a linear fit to the MSD of the center of mass of the FUS-LCD chains, we obtained a rough estimate of the translational diffusion coefficient of 0.26 ·10*^−^*^3^ nm^2^/ns (Figure S2), which is within the range of experimental values reported by Fawzi and coworkers (0.17 to 0.4 ·10*^−^*^3^ nm^2^/ns).^24,25^

Figure 1A shows the MSD, averaged over all individual backbone C*_α_*-atoms in the protein chains, plotted as a function of the lag time. In the condensate, an average translational slowdown relative to the dilute phase of up to two orders of magnitude can be observed (the black dashed line in Figure 1A represents the mean MSD of the C*_α_*-atoms in the condensate multiplied by 100). This result is also in line with previous simulations showing strongly retarded overall diffusion of proteins in crowded environments.^57–60^ To analyze this slowdown in further detail, MSDs of groups of increasing numbers of consecutive C*_α_*-atoms in the FUS-LCD chain were calculated, with group sizes ranging from a single C*_α_* atom to the whole chain (163 C*_α_*-atoms)

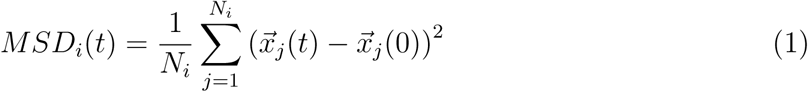

**Figure 1:**
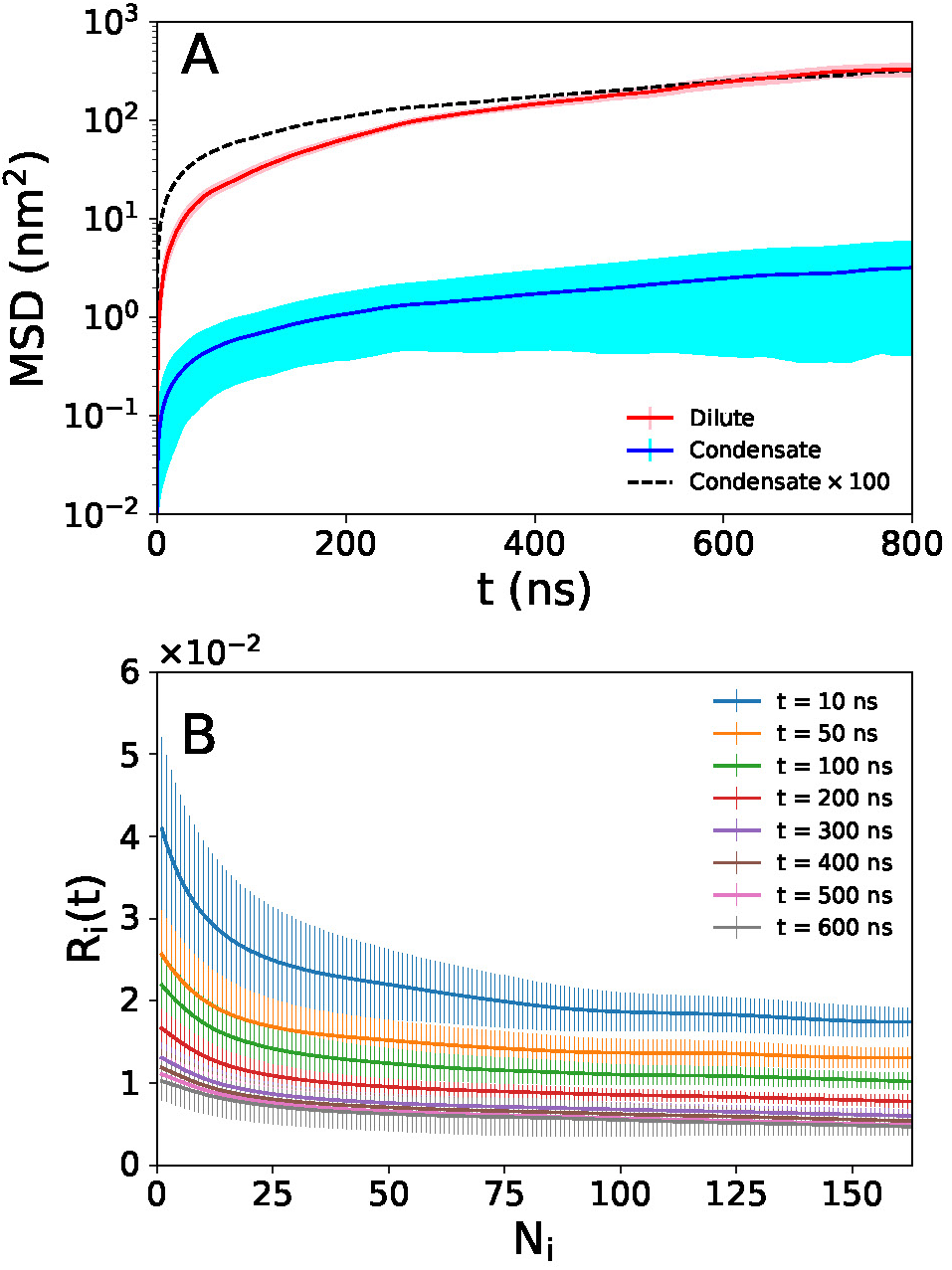
A) Mean squared displacements of C*_α_*-atoms in the protein backbones in the dilute (red) and condensate (blue) systems plotted as a function of the lag time t. The solid lines represent the mean MSD and the error bars denote the standard deviations over all the C*_α_*-atoms considered. The black dashed line represents the condensate MSD multiplied by 100. B) Ratio of the mean MSDs of C*_α_*-atoms in the condensate and the dilute systems at selected lag times (10, 50, 100, 200, 300, 400, 500, 600 ns) plotted against the number of C*_α_*-atoms in a group. The error bars denote the standard deviations over the 8 protein chains in the simulation system.

In equation 1, 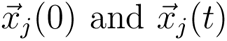 denote the position vectors of the j*^th^* C*_α_* atom at a time origin and at a lag time t later, respectively. Each group of size N*_i_* (the number of C*_α_*-atoms in the i*^th^* group) has 163 − N*_i_* + 1 instances or subgroups, and the MSD was calculated by averaging over all subgroups.

Figure 1B shows the ratio of MSDs in the condensate and the dilute system as a function of N*_i_* at selected lag times

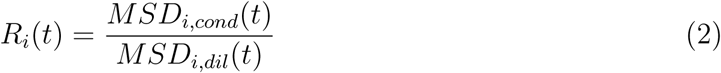

A small value of R*_i_*(t) indicates a strong slowdown of dynamics in the condensate phase. As shown in Figure 1B, the dynamical retardation is more pronounced at longer lag times compared to shorter ones. Additionally, the slowdown is stronger for larger groups of atoms, whereas smaller groups of atoms at shorter lag times experience the least dynamical retardation. The differences in R*_i_*(t) diminish for longer lag times, indicating that the retardation as a function of lag time is nonlinear and that beyond a certain point, the dynamics are similarly slowed down.

Our findings show that motions of larger groups of atoms on larger length-scales (as expected at longer times) through the viscous condensate environment are subject to stronger retardation. This is in line with the theoretical expectation that the friction experienced by a (spherical) probe particle upon moving through a macromolecular environment is reduced due to an entropic depletion effect,^61^ leading to an effective viscosity that depends on the size of the diffusing molecule relative to the mesh spacing of the surrounding macromolecular network.^33,62^

The above analysis provides a measure of the retardation of the translational motions of the protein backbone in the condensate. To more closely understand this dynamic slowdown, the reorientation motions of the protein backbone were analyzed separately from the side-chains. To this end, the reorientation of the vectors between two C*_α_*-atoms at varying sequence separations were studied. First, the orientational autocorrelation functions (ACFs)

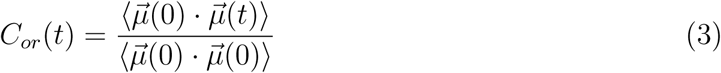

were calculated for the C*_α_* −C*_α_* vectors 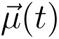 at different separations (N*_Cα−Cα_* = 1 to 162), averaged over all possible instances of N*_Cα−Cα_* . The brackets ⟨. . .⟩ indicate the average over all time step differences t. The ACF in equation 3 describes the reorientation motion of the C*_α_* − C*_α_* vectors via the angle θ*_t_* formed by a vector during the time interval t. The mean ACFs were fitted to biexponential functions (Eq. 4) and the mean orientational time-scales (Eq. 5) were obtained by integration

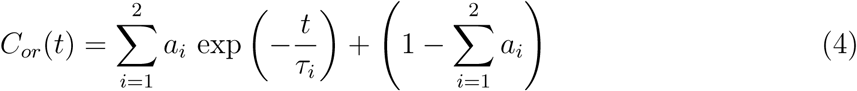

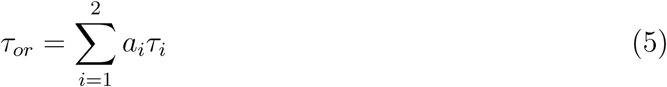

The amplitudes a*_i_* were normalized to obtain relaxations between 1 and 0. ACFs for selected C*_α_* − C*_α_* separations are shown in Figure S3. The applicability of the biexponential fit was decided by calculating the R^2^ (coefficient of determination) values at each separation and comparing mono-, bi-, and triexponential fits (Figure S4). The triexponential fits were not superior to the biexponential ones, so the biexponentials were used.

The same approach was taken to quantify the magnitude of distance fluctuations of the

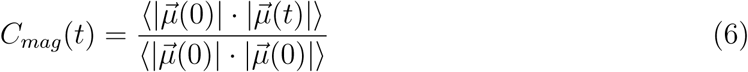

where 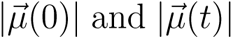 are the magnitude of the vector at time origin and at a lag time t, respectively. Also in this case, biexponential functions accurately described the relaxations (Figures S5 and S6).

Figures 2A and 2B show the orientational time-scales for the dilute 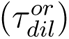 and the condensate 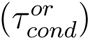 systems, respectively. The ratio of the two time constants is shown in Figure 2C. Figures 2D, 2E, and 2F show the analogous graphs for the C*_α_* − C*_α_* distance fluctuations. In this case the correlation times are denoted by 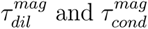 for the dilute and the condensate phase, respectively.

**Figure 2:**
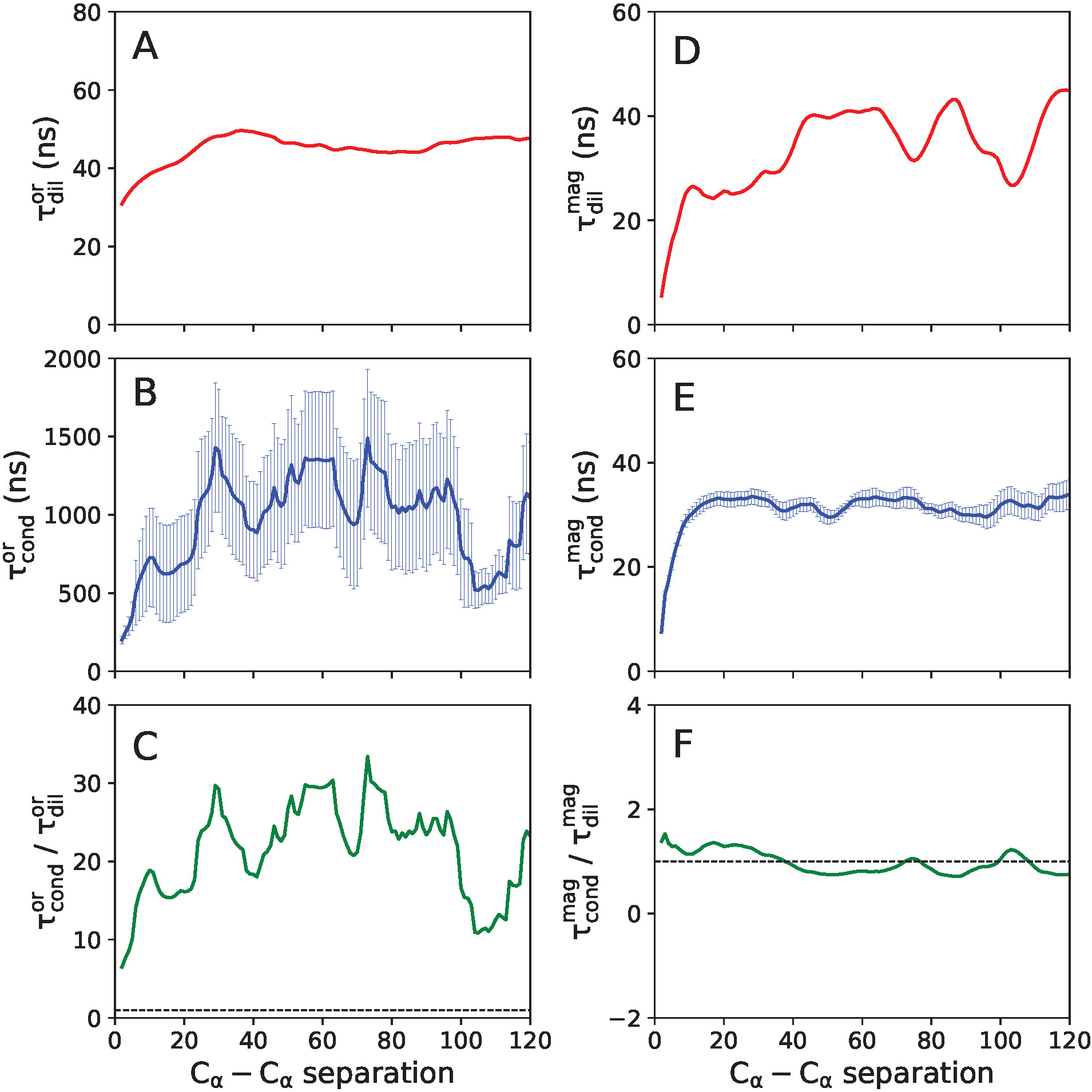
Protein backbone dynamics for FUS-LCD chains. A) and B) represent the time-scales of orientational dynamics as a function of C*_α_* − C*_α_* separation for the dilute and the condensate systems respectively. C) represents the ratio of these two time-scales. D), E), and F) show the analogous properties for the C*_α_* − C*_α_* distance fluctuations. The error bars denote the standard errors over the 8 proteins in the condensate system.

Figure 2A shows that retardation of orientational backbone dynamics is observed with an increase in N*_Cα−Cα_* . For larger length-scales (sequence separation N*_Cα−Cα_* > 40), the value of 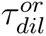 plateaus. A qualitatively similar behavior is also observed for 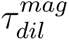 (Figure 2D).

A smooth N*_Cα−Cα_* dependence is found for 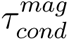 (Figure 2E), while 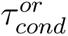 (Figure 2B) shows N*_Cα−Cα_* larger variations.

The time-scale ratio depicted in Figure 2C illustrates that the reorientation motions of the backbone are slowed down by factors of approximately 5-15 (about one order of magnitude) in the condensate compared to the dilute system. In contrast, the C*_α_* −C*_α_* distance dynamics exhibit no noticeable slowdown in the condensate system (Figure 2F). This apparent contradiction can be resolved by considering the nature of dynamical variables analyzed. While the reorientation dynamics of the protein backbone involves motions of backbone atoms through a dense and viscous environment, the distance magnitude fluctuations pertain to the local motions of the C*_α_*-atoms around their average positions. Thus, in the context of the above results on the retardation of global protein translational dynamics, we conclude that protein dynamics involving large-scale motions are more strongly affected by the condensate environment.

Next, the motions of the amino acid side-chains were investigated in terms of rotamer jumps. The side-chain rotamer dynamics were analyzed by computing the normalized time ACFs C*_χ_*(t) of the dihedral angles χ*_i_* (i = 1, 2, 3, depending on the length of the side-chain)

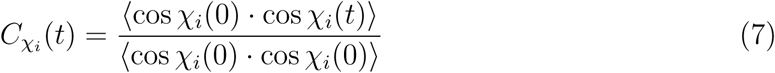

For each amino acid occurring in the FUS-LCD sequence that at least has the χ_1_ dihedral (Asn, Asp, Gln, Ser, Thr, and Tyr), for each χ*_i_*, the ACFs (Eq. 7) were averaged over all the side-chains present in the system. These mean ACFs were fitted by biexponential functions (Eq. 4) and integrated (Eq. 5) to obtain the correlation times, which are plotted in Figure 3. The individual ACFs are shown in Figures S7 and S8. Biexponential functions accurately describe the ACFs (Figure S9).

**Figure 3:**
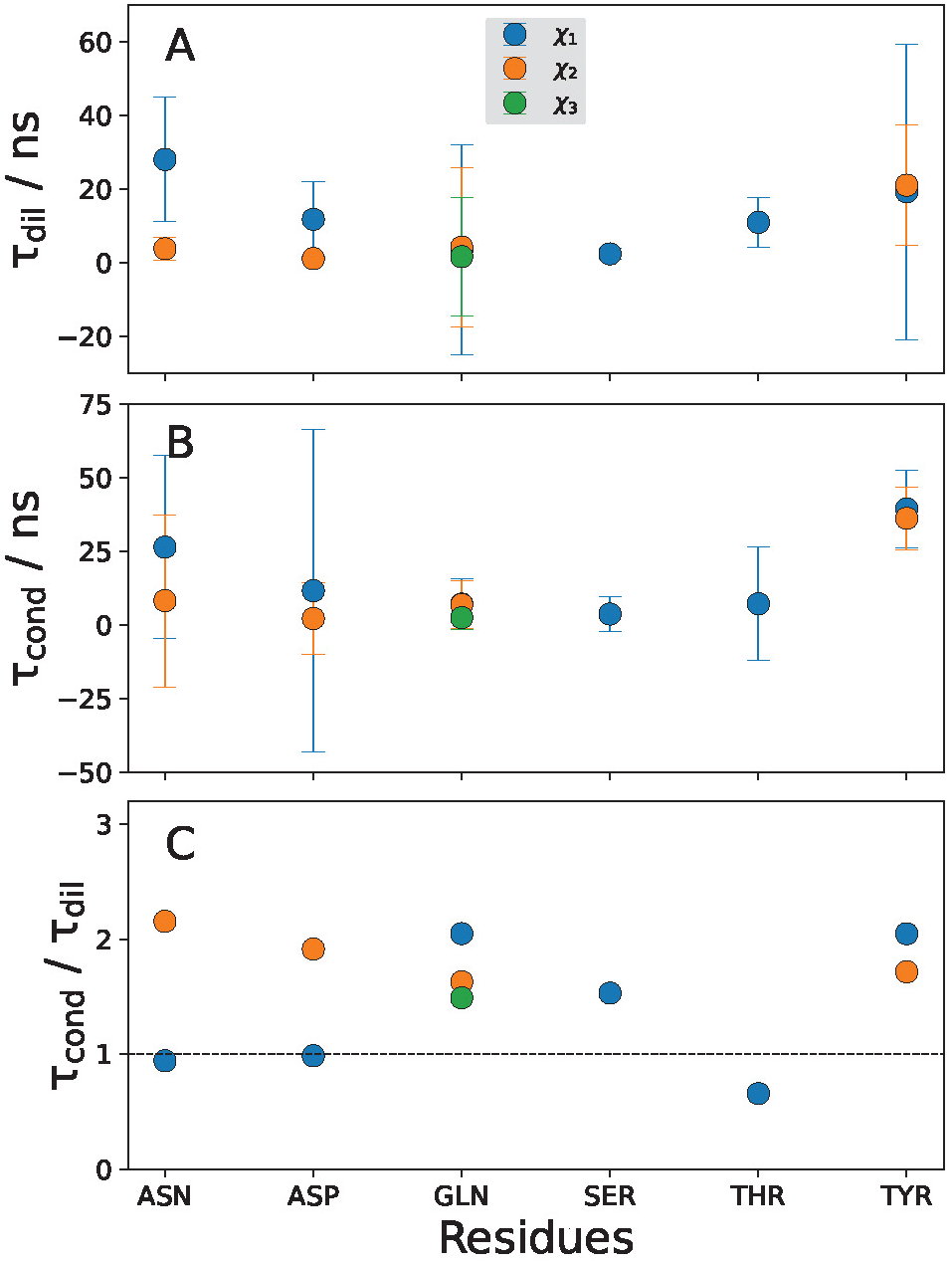
time-scales of side-chain dihedral angle rotations in the dilute (A) and the condensate (B) systems. C) shows the ratio of the time constants in the two systems. The χ_1_, χ_2_, and χ_3_ dihedral angles of the respective residues are plotted in blue, orange, and green, respectively. The error bars denote the standard errors over all the respective residues present in the system.

Figures 3A and 3B present the time-scales in the dilute (τ*_dil_*) and the condensate (τ*_cond_*) systems, respectively. The ratio of τ*_cond_* and τ*_dil_* is shown in Figure 3C. The side-chain dynamics are not significantly slowed in the condensate. At first sight, it may seem surprising that rotamer jumps are not affected even for the tyrosine side-chains, which are known to play a distinct role for mediating residue-residue interactions ins FUS condensates. ^47^ Averaged over all side-chains and dihedral angles (χ_1_, χ_2_, χ_3_), the retardation is about twofold. This small change is similar to the backbone C*_α_* − C*_α_* magnitude fluctuations, supporting the notion that the time-scales of the small-amplitude motions are not much affected in the condensate. As described above, at short length-scales the entropic depletion effect reduces the effective viscosity of the macromolecular network, and hence fast local motions do not experience a strong retardation.

Finally, we aimed at obtaining a more complete picture and analyzed how the heterogeneous retardation of the protein dynamics in the condensate described above influences the dynamics of the surrounding water molecules. As protein and water motions are coupled,^63–70^ and since water motions on a local scale are fast (typically in the picoseconds regime), we expected them to couple most strongly to the fast local small-amplitude protein motions, and thus to find a similar modest condensate-related slowdown as we found for the local protein motions, i.e., about twofold. To test that hypothesis, we probed the reorientation motions (rotations) of the water molecules in the protein hydration layer (PHL), which was defined as the region within 0.5 nm of the protein surface. To quantify the time-scales of water rotations, we computed the time ACFs of the dipole moment vector of each water molecule, tracking only those water molecules that remained within the PHL for the duration of the analysis. In this case, µ⃗(t) in equation 3 denotes the normalized dipole moment vector of a single water molecule (for a rigid water model, as used in this work, an O-H bond vector (and not the dipole moment) can be used for the analysis). As described above, the ACFs were fitted to biexponential functions and integrated to obtain the rotational correlation times. The distributions obtained for the dilute and condensate phases are plotted in Figure 4A.

**Figure 4:**
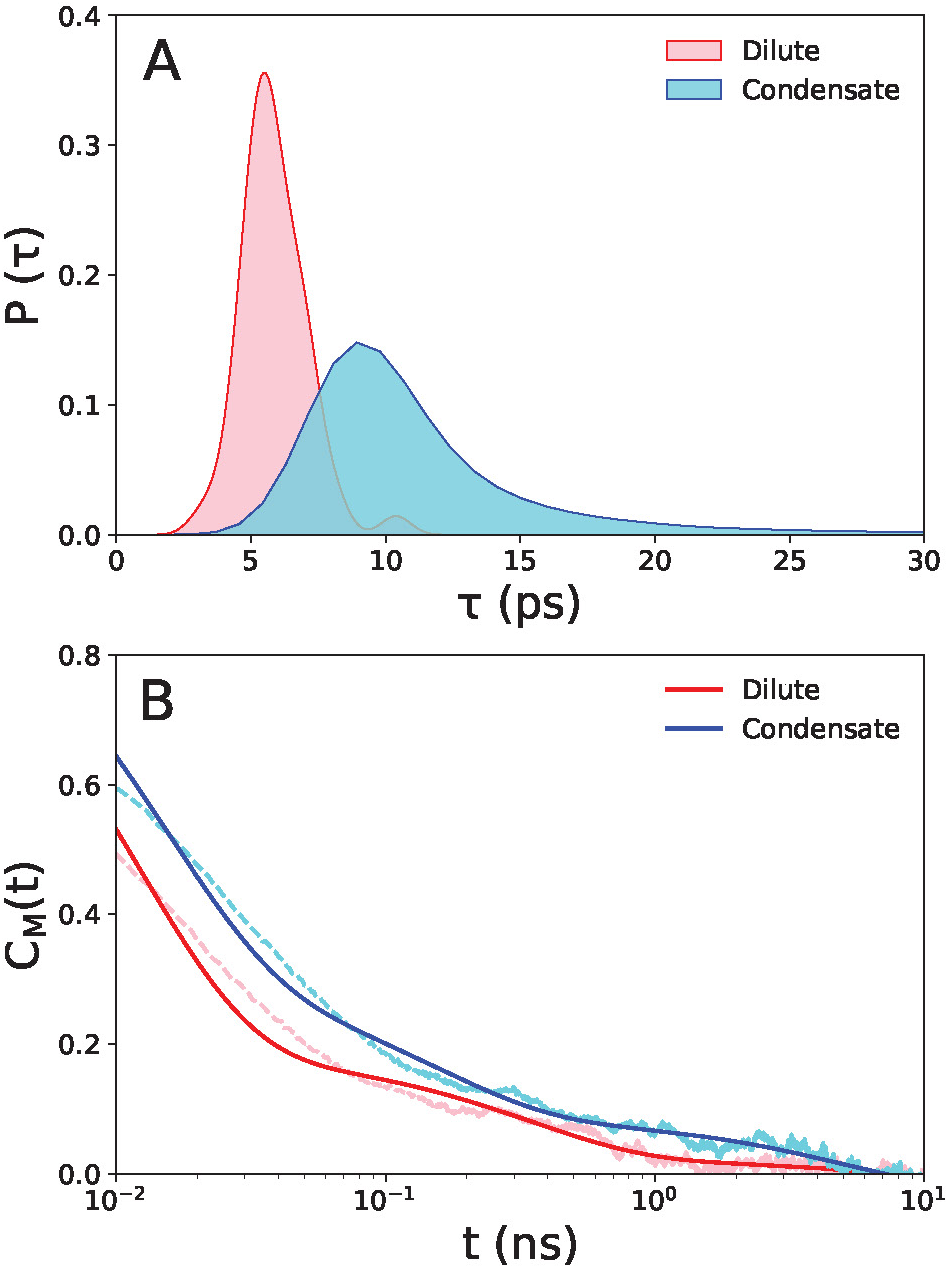
Dynamics of protein hydration layer (PHL) in the dilute (red) and condensate (blue) phases. The PHL is defined by a cut-off distance of 0.5 nm from the protein. A) Distributions of rotational correlation times. B) ACFs of total dipole moment (and triexponential fits, solid lines) of PHL water molecules in the dilute and condensate phases plotted on a semi-log scale.

Figure 4A shows that the time-scale distribution for the condensate system is broader and slightly shifted compared to the dilute phase, indicating a partial slowdown in the rotational dynamics of water in the PHL in the condensate. The mean correlation times for the dilute and the condensate systems are 6 ps and 12 ps, respectively. Hence, the water reorientations in the condensate are slowed down by a factor of two.

While the above analysis provides a measure of the dynamical slowdown at the level of single water molecules, a more global picture can be obtained by comparing the collective relaxation of the entire protein hydration layer, as described by the ACF of the total dipole moment C*_M_* (t)

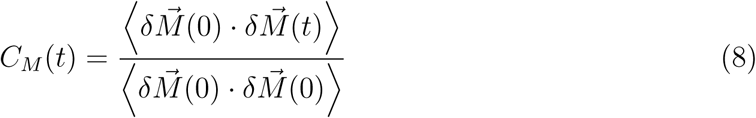

In equation 8, the total dipole moment fluctuation is denoted by 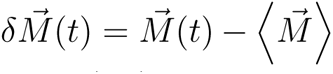,

where 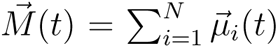 is the total dipole moment vector of the PHL, 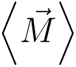 denotes the time-averaged dipole moment, and 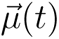 is the dipole moment of the i*^th^* water molecule in the PHL. The C*_M_* (t) of the dilute (red) and the condensate (blue) phases are plotted in Figure 4B. The dashed lines represent the raw ACFs, whereas the solid lines denote their fits. In this case, a triexponential function provided better fits 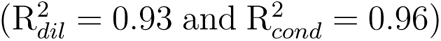 than a biexponential function 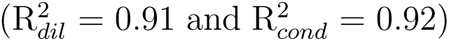. The average correlation times thus obtained are 101 ps and 670 ps for the dilute and the condensate systems, respectively. The hydration water analysis thus shows that single-molecule rotational water motions are slowed down twofold, while overall collective dynamics of the ensemble of hydration water molecules are slower and affected to a greater extent by the condensate environment. These results are in qualitative agreement with recent vibrational spectroscopy studies,^71,72^ which showed that local vibrational probes, such as the amide backbone C=O stretch or the arginine side-chain C-N stretches, are slowed modestly in the condensate environment compared to the dilute phase (about 2-6 fold, in agreement with our above results, e.g., of the the local side-chain motions). The dynamic retardation of these local protein modes is transmitted to the surrounding water in the PHL via the H-bond network.

The interpretation of the relevance of the (overall only relatively modest) retardation of the water dynamics in the condensate phase in the context of biological function is challenging. However, we would like to point out that slow water dynamics and a concomitant stiffening of the water H-bond network are generally associated with lower water entropy.^73–77^ The associated solvation-related contributions to the thermodynamic driving forces of condensate formation can be sizeable in magnitude, especially when the water molecules are constrained by the strongly confined environment in the condensate.^78,79^ However, even at protein concentrations as high as 300 mg/ml and beyond, the water volume fraction in the condensate is still relatively large (above 60%), ensuring that the condensates retain a sufficient liquid-like character to maintain a high degree of biomolecular dynamics.

In conclusion, in this work we used atomistic MD simulations to probe the dynamic slowdown of an intrinsically disordered protein and the water in its hydration layer in bio-condensates, using one-component FUS-LCD condensates as a model system. The crowded milieu in the dense condensate phase slows down both protein and water dynamics, but to markedly different extents depending on the length-scale of the motions involved. The retardation of dynamics is non-universal and heterogeneous, with dynamic modes that involve large-scale motions being affected more strongly than small-scale local fluctuations. The most significant dynamical retardation in the condensate of up to almost two orders of magnitude is found for the translational motions of the protein backbone. The slowdown increases with increasing number of atoms involved in the motion, that is, collective large-amplitude motions are affected the most. For small-amplitude motions such as C*_α_* distance fluctuations and side-chain dihedral rotations, the dynamical slowdown by the condensate is not very pronounced. However, the orientational dynamics of C*_α_* −C*_α_* vectors, which require movement of the protein backbone chain through the condensate interior, are found to be slowed down by one order of magnitude.

Taken together, this work provides detailed microscopic insights into how slow translational diffusion on a global scale is linked to fast local structural dynamics. The latter might be important for the biological function of FUS, for example by fostering molecular recognition processes such as FUS binding to RNA or to protein factors involved with transcription initiation and regulation. The connection found here between the dynamic slowdown and the time- and length-scales of the associated protein motions might be transferable to other biocondensates and thus have more general implications beyond the specific system studied in this work. The precise link between protein dynamics and biological function is very case-specific, but it has been shown for many biomolecular systems that dynamics are required for function. It is therefore essential for the proteins to retain their dynamic personalities ^80^ in dense biocondensates, as exemplified in this work.

## Acknowledgements

This project received funding from the European Union’s Horizon 2020 research and innovation programme under the Marie Sk-lodowska-Curie grant agreement No 801459 - FP-RESOMUS and from the Deutsche Forschungsgemeinschaft (DFG) under Germany’s Excellence Strategy - EXC 2033 - 390677874 - RESOLV.

## Supporting Information Available

Methods (system set-up and simulation details); MSD over lag time plot and linear fit of translational diffusion coefficient; Selected ACFs for C*_α_* − C*_α_* reorientation dynamics for selected separations in the condensate system; R^2^ values plotted against C*_α_* − C*_α_* separations for reorientation dynamics in dilute and condensate systems for different fitting functions; Selected ACFs for C*_α_* − C*_α_* distance fluctuations for selected separations in condensate system; R^2^ values plotted against C*_α_*− C*_α_* separations for distance fluctuations in dilute and condensate systems for different fitting functions; ACFs of side-chain rotamer dynamics in dilute and condensate systems; R^2^ values for different fitting functions in the side-chains dynamics ACFs.

## Supporting Information

### Methods

#### System preparation

The intrinsically disordered low complexity domain (LCD) of the human Fused in Sarcoma (FUS) RNA-binding protein (UniProt ID: P3563, residues 1 to 163 of a total 526 residues) was simulated in this study. The initial atomic coordinates of FUS-LCD were generated using AlphaFold, ^S1,S2^ which predicted an extended conformation without any secondary structure.

As explained in detail in our previous work,^S3^ the initial equilibrated condensate system at a protein concentration of 350 mg ml*^−^*^1^ was obtained from coarse-grained (CG) MD simulations with the Martini 2.2^S4^ force field. Scaled protein-protein Lennard-Jones (LJ) interactions were used to maintain the liquid-like dynamical features of the condensate, as suggested by Benayad et al. ^S5^ Eight copies of the coarse-grained FUS-LCD chain were inserted in a cubic periodic simulation box along with 4366 CG water beads (1 Martini water bead corresponds to 4 atomistic water molecules) and 16 sodium ions to neutralize the net charge of the simulation box.

The system was energy minimized and simulated for 1 µs under NpT conditions at T = 300 K and p = 1 bar. The resulting box dimensions were 8.9 nm. Temperature and pressure were maintained using a weak coupling thermostat and barostat,^S6^ respectively. The coarse-grained simulations were performed using the recommended ”new rf” settings,^S7^ which included the use of a 1.1 nm cut-off for the non-bonded interactions and a 20 fs time step to integrate the equations of motion.

The CG simulations enabled fast equilibration of the protein configurations. However, realistic dynamical data (time scales) can not be obtained from this approach. Hence, the CG simulations were only used for initial equilibration but not for analysis. All production simulations were performed with the all-atom force field amber99SB-disp.^S8^ First, the systems obtained from CG equilibration were back-mapped to the all-atom level using the initram.sh and backward.py scripts.^S9^ The resulting atomistic proteins were solvated with 16509 water molecules. 79 sodium and 63 chloride ions were added to neutralize the charges from the protein chains and to have a salt concentration of 150 mM, corresponding to physiological ion concentrations. The protein, water and ions were described by the a99SB-disp force field (and the corresponding 4-point water model derived from TIP4P-D),^S8^ which was shown to be a good choice for FUS in a recent study.^S10^

A reference dilute system was also simulated. It included a single FUS-LCD chain in 45681 water molecules along with 134 sodium and 132 chloride ions in a 11.2 nm cubic periodic box. The salt concentration in this case was also 150 mM. Snapshots of the dilute and condensate systems are shown in Figs. S1A and S1B respectively.

**Figure S1:**
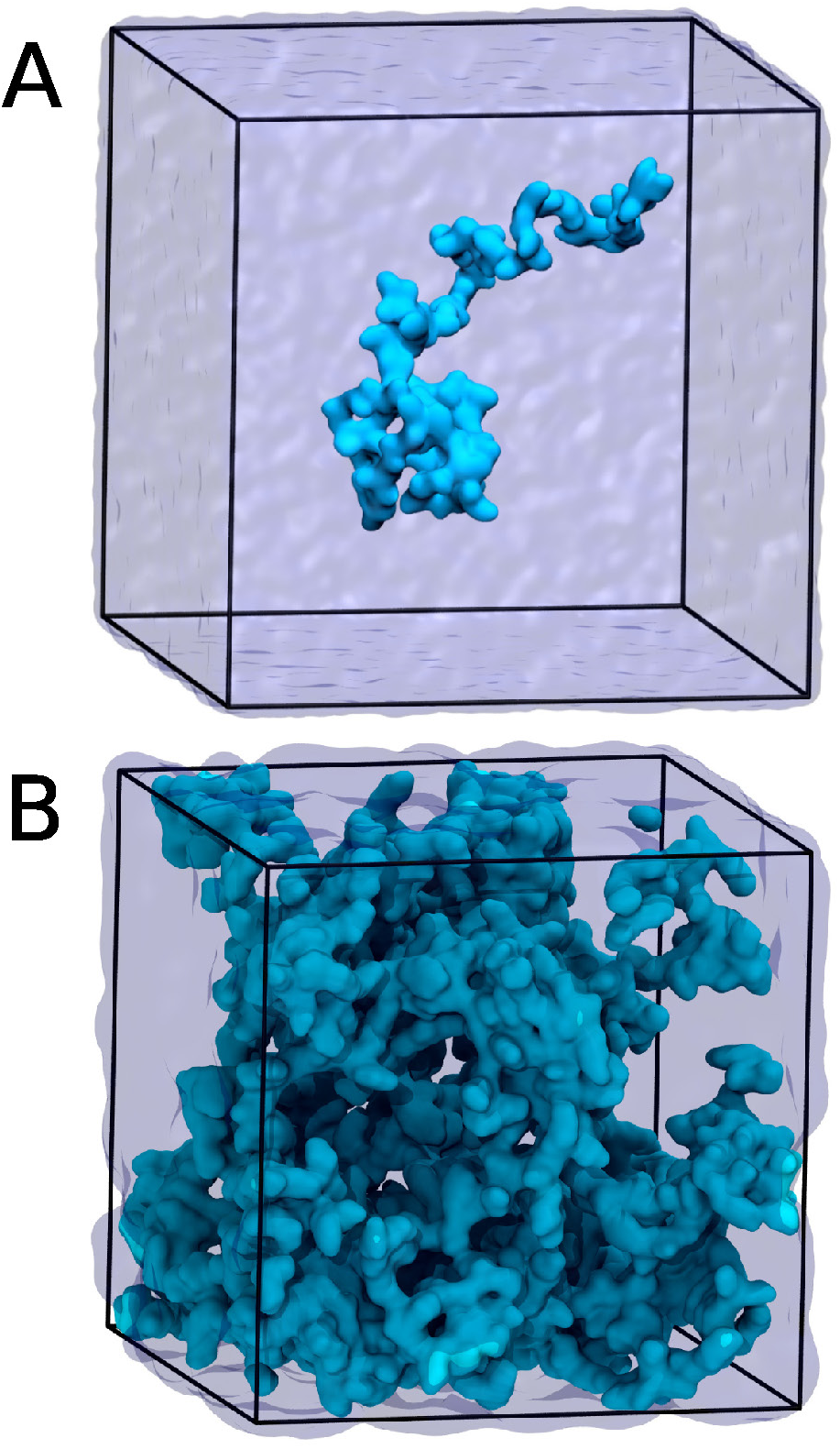
Snapshots of the equilibrated simulation boxes of (A) dilute and (B) condensate systems.

### Simulation details

All simulations were performed using the GROMACS (version 2021.1) molecular dynamics simulation package.^S11^ Both the systems were first energy minimized using the steepest descent algorithm and then equilibrated with harmonic position restraints on the protein backbone for 10 ns (restraining force constants of 1000 kJ mol*^−^*^1^ nm*^−^*^2^) under NpT conditions at T = 300 K and p = 1 bar. This was followed by further NpT equilibration under the same conditions for 100 ns without any position restraints. Thereafter, the systems were equilibrated for 100 ns at constant volume, followed by the final production runs for 1 µs in the NVT ensemble. Coordinates were saved to disk every 10 ps. For the analysis of water dynamics, 100 ns production runs were performed with a data saved to disk every 1 ps. In these simulations, the velocity rescaling thermostat with a stochastic term^S12^ and the Berendsen barostat^S6^ were used to control temperature and pressure, respectively.

All atomistic MD simulations were done using the leapfrog integrator with a time step of 2 fs. Protein bonds and internal degrees of water molecules were constrained using the LINCS and SETTLE algorithms, respectively. Short-range Coulomb and Lennard-Jones interactions were calculated up to an interparticle distance cut-off of 1.0 nm. Long-range electrostatic interactions were treated with the particle-mesh Ewald method ^S13^ with a grid spacing of 0.12 nm and cubic spline interpolation.

**Figure S2:**
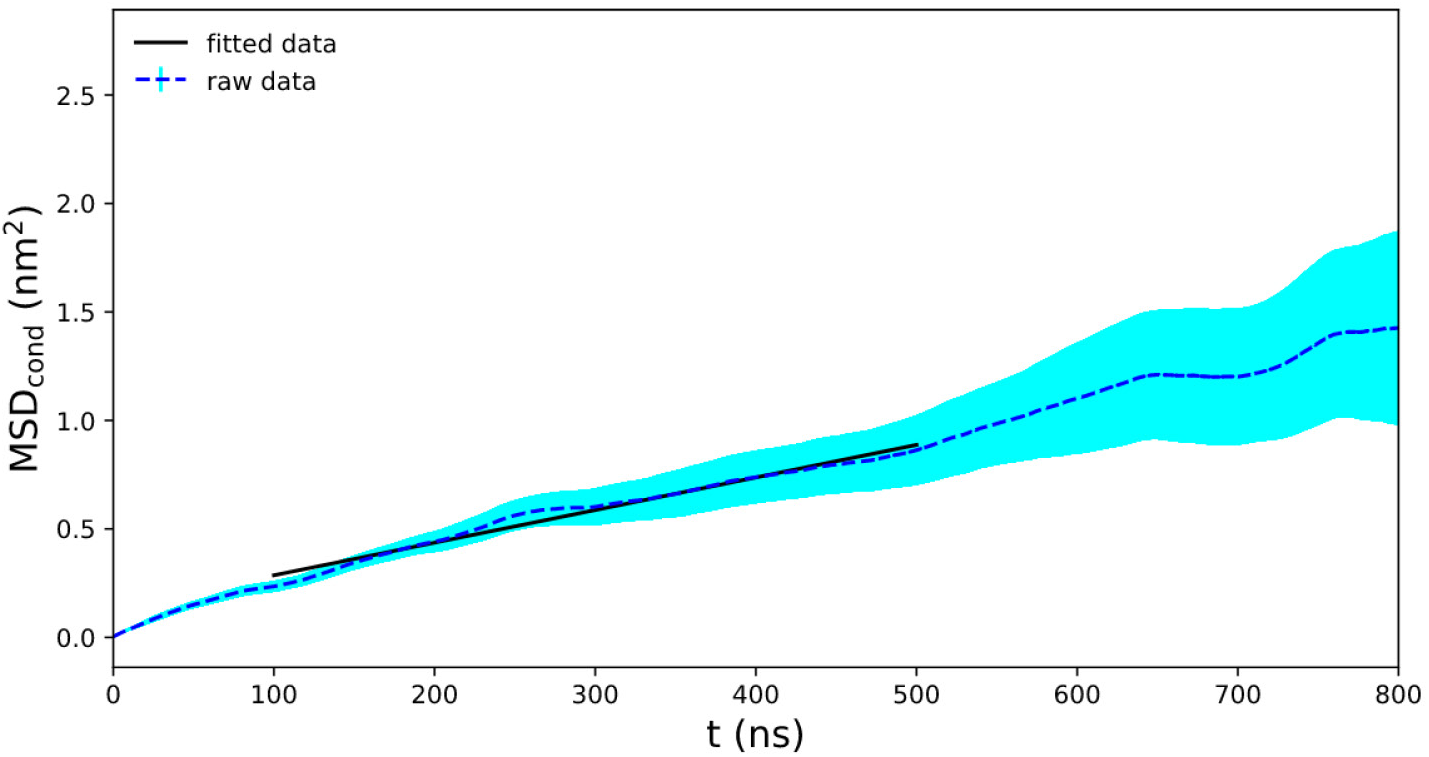
Mean squared displacement (MSD) of the centers of mass of the FUS-LCD chains in the condensate simulation system. The dashed line represents the raw data, averaged over the eight proteins in the simulation box. The cyan shaded area shows the standard deviation over the eight proteins. The solid line depicts the linear fit from 100 ns to 500 ns; the slope is 6D, with D the apparent translational diffusion coefficient. The obtained value of D = 0.26 · 10*^−^*^3^ nm^2^/ns is within the range of values from FRAP and NMR experiments reported by Fawzi and coworkers^S14,S15^ (0.17 to 0.4 ·10*^−^*^3^ nm^2^/ns). However, we consider it to be only a rather rough estimate, because the fit assumes a linear relationship between MSD and lag time and thus neglects subdiffusive behavior, which is expected to play a role for the confined motions in the condensate. Furthermore, we did not correct for finite box size effects^S16^ because the viscosity of the simulation box is unknown. Such correction would increase the value slightly.

**Figure S3:**
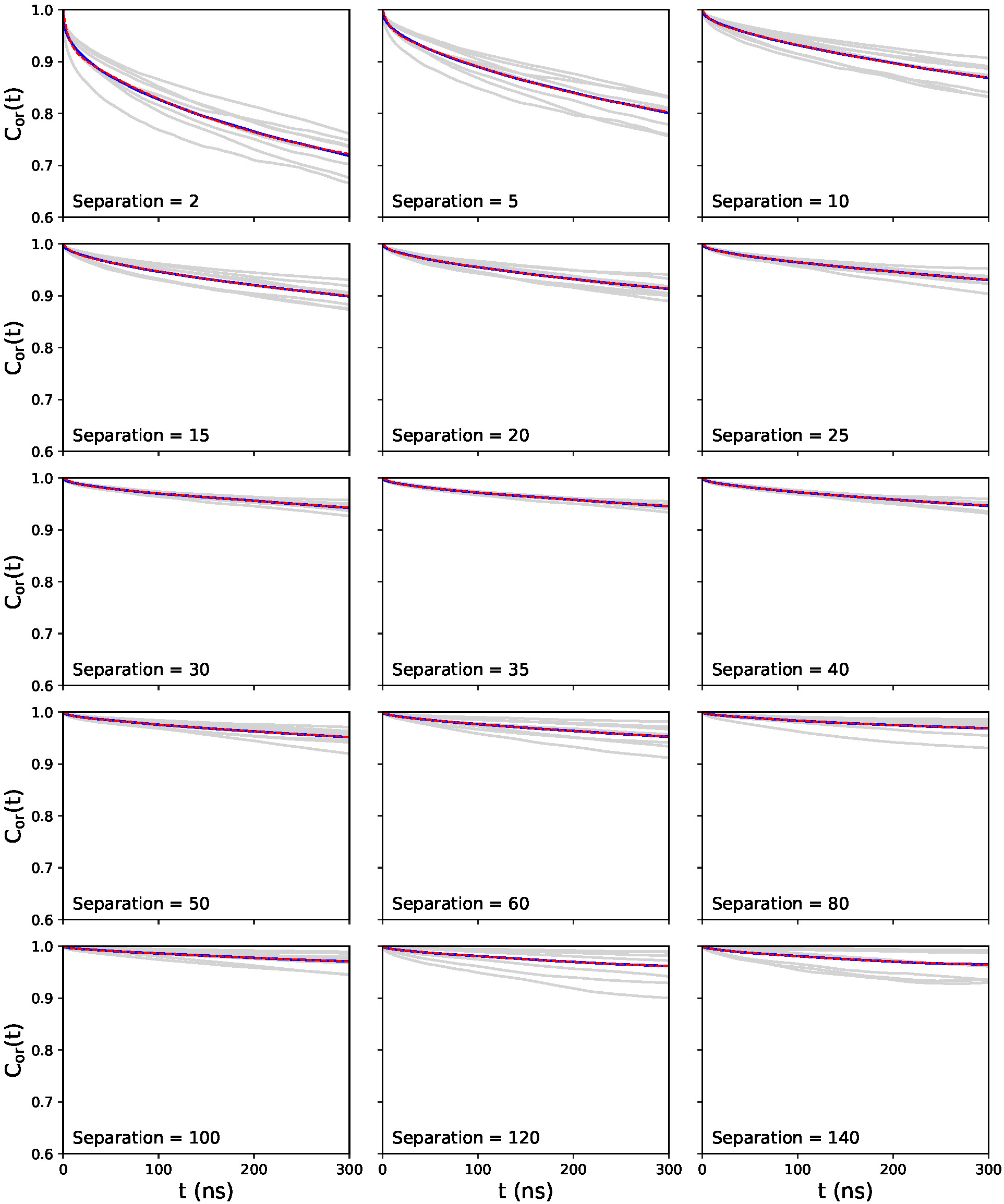
Orientational autocorrelation functions of the C*_α_*−C*_α_* vectors at selected sequence separations in the condensate system. The light grey curves represent the 8 individual proteins in the system. The bold blue line is the average ACF over the 8 individual ones. The dashed red lines show the biexponential fits.

**Figure S4:**
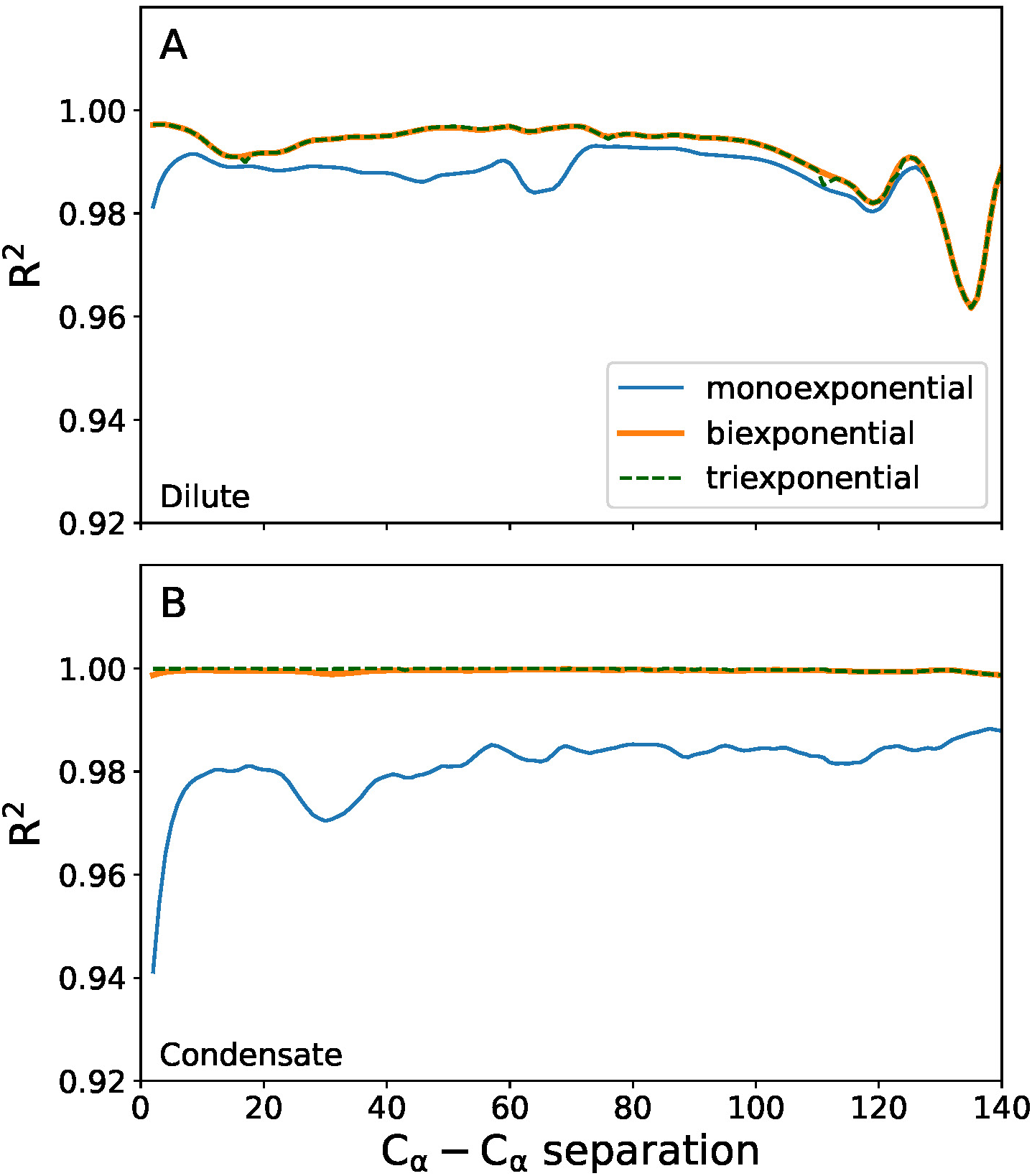
Coefficient of determination (R^2^) for mono-, bi-, and triexponential fits of the ACFs of C*_α_* − C*_α_* vector orientation dynamics as a function of C*_α_* − C*_α_* separation in (A) dilute and (B) condensate systems.

**Figure S5:**
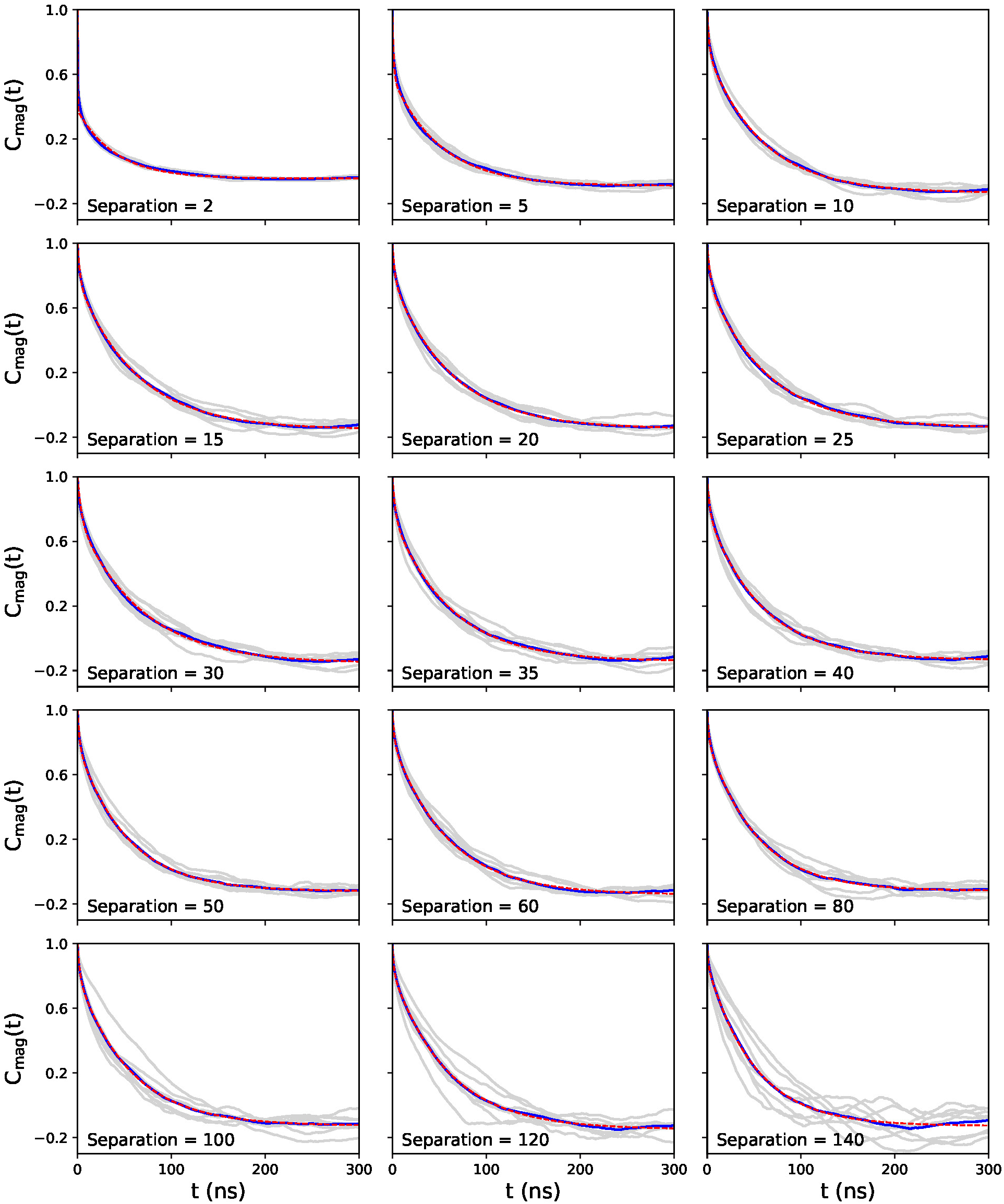
ACFs of the C*_α_* − C*_α_* distance fluctuations at selected sequence separations in the condensate system. The light grey curves represent the 8 individual proteins in the system. The bold blue line is the average ACF over the 8 individual ones. The dashed red line denotes the biexponential fit.

**Figure S6:**
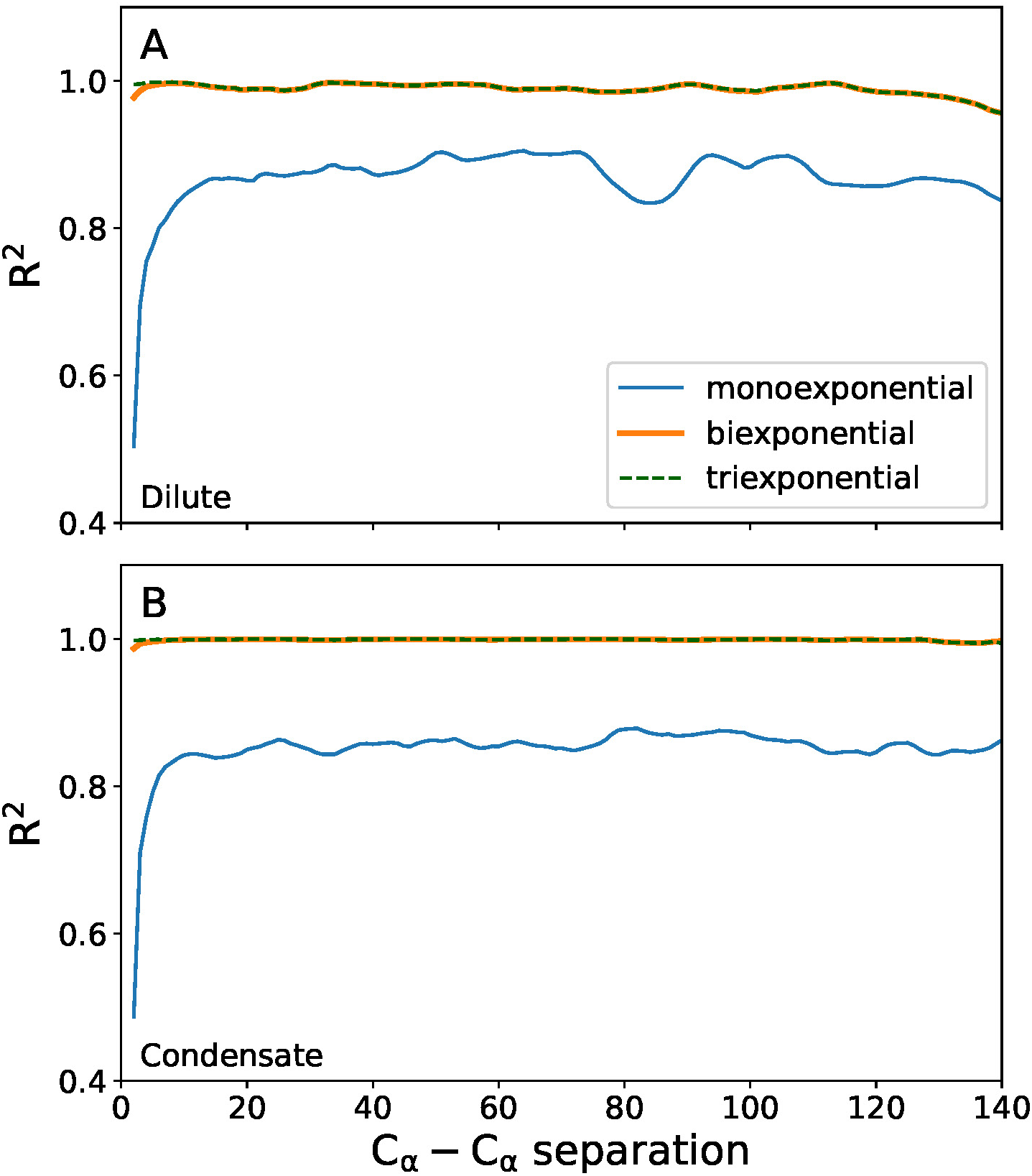
Coefficient of determination (R^2^) for mono-, bi-, and triexponential fits of the ACFs of C*_α_* − C*_α_* distance fluctuations as a function of C*_α_* − C*_α_* separation in (A) dilute and (B) condensate systems.

**Figure S7:**
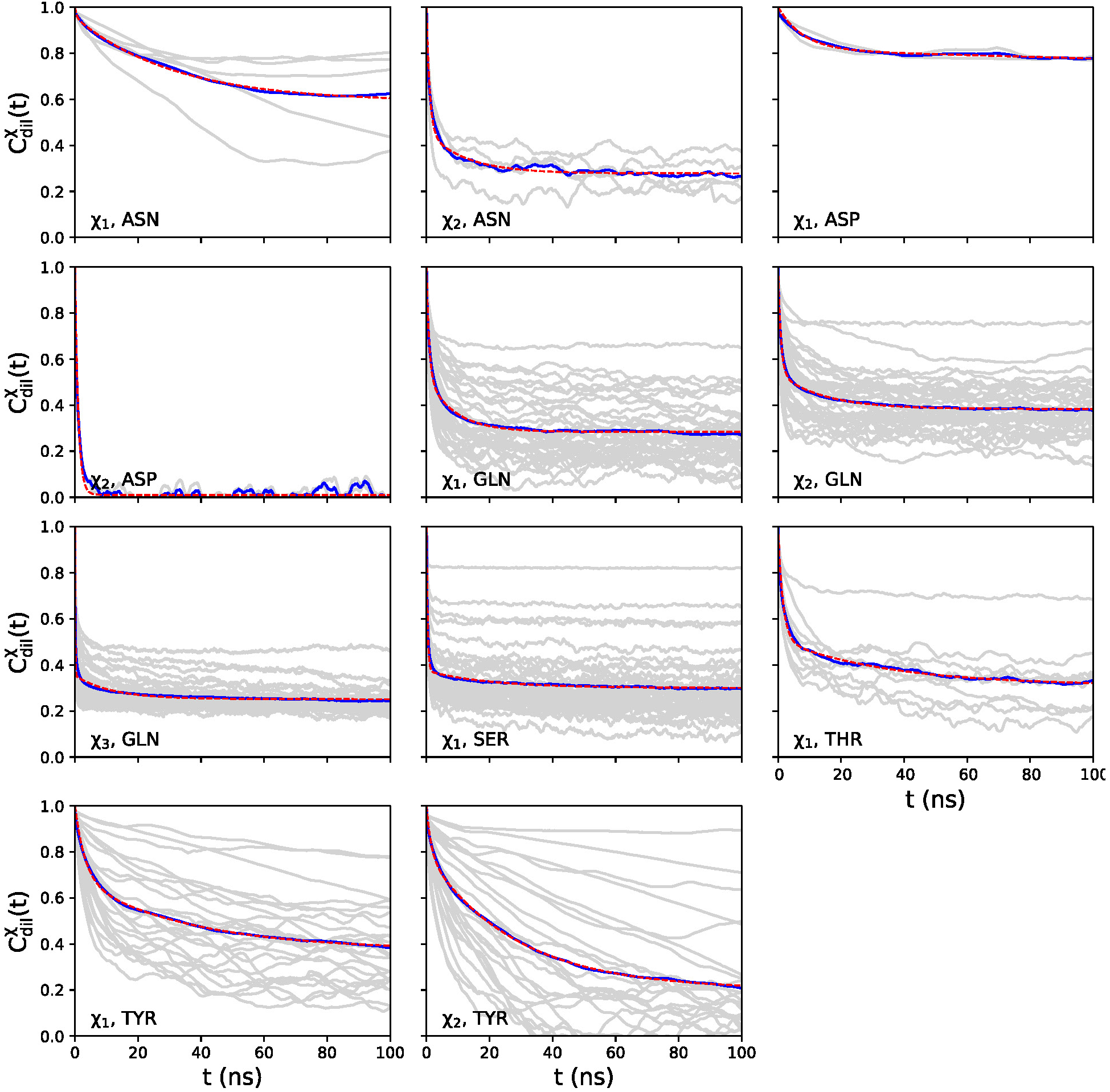
Side-chain dihedral autocorrelation functions of the protein in the dilute system. The light grey curves represent the individual side-chains of the same type present in the system. The bold blue line is the average ACF. The dashed red line denotes the biexponential fit to the mean ACF.

**Figure S8:**
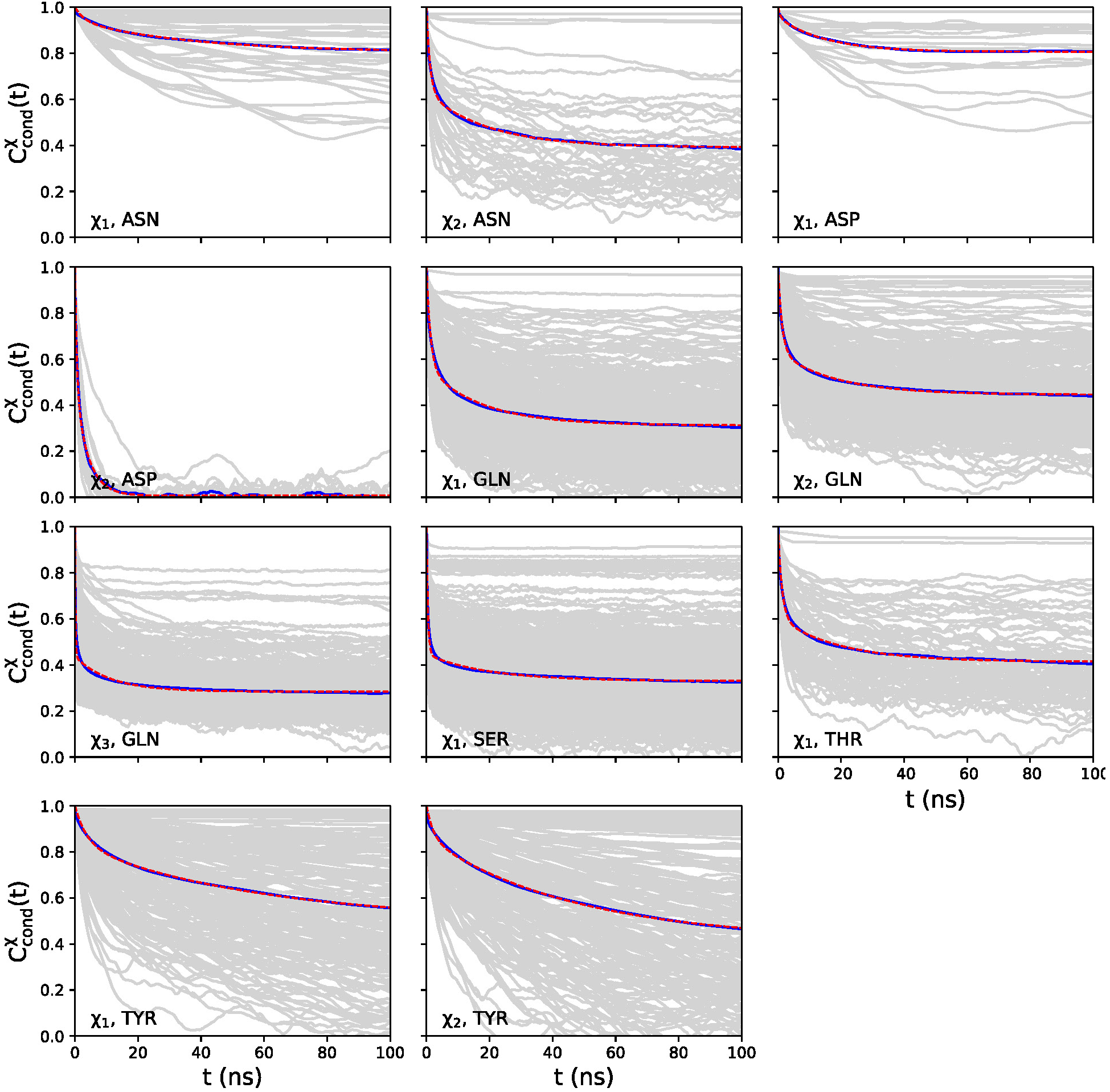
Side-chain dihedral ACFs in the condensate system. The light grey curves represent the individual side-chains of the same type present in the system. The bold blue line is the average ACF. The dashed red line denotes the biexponential fit.

**Figure S9:**
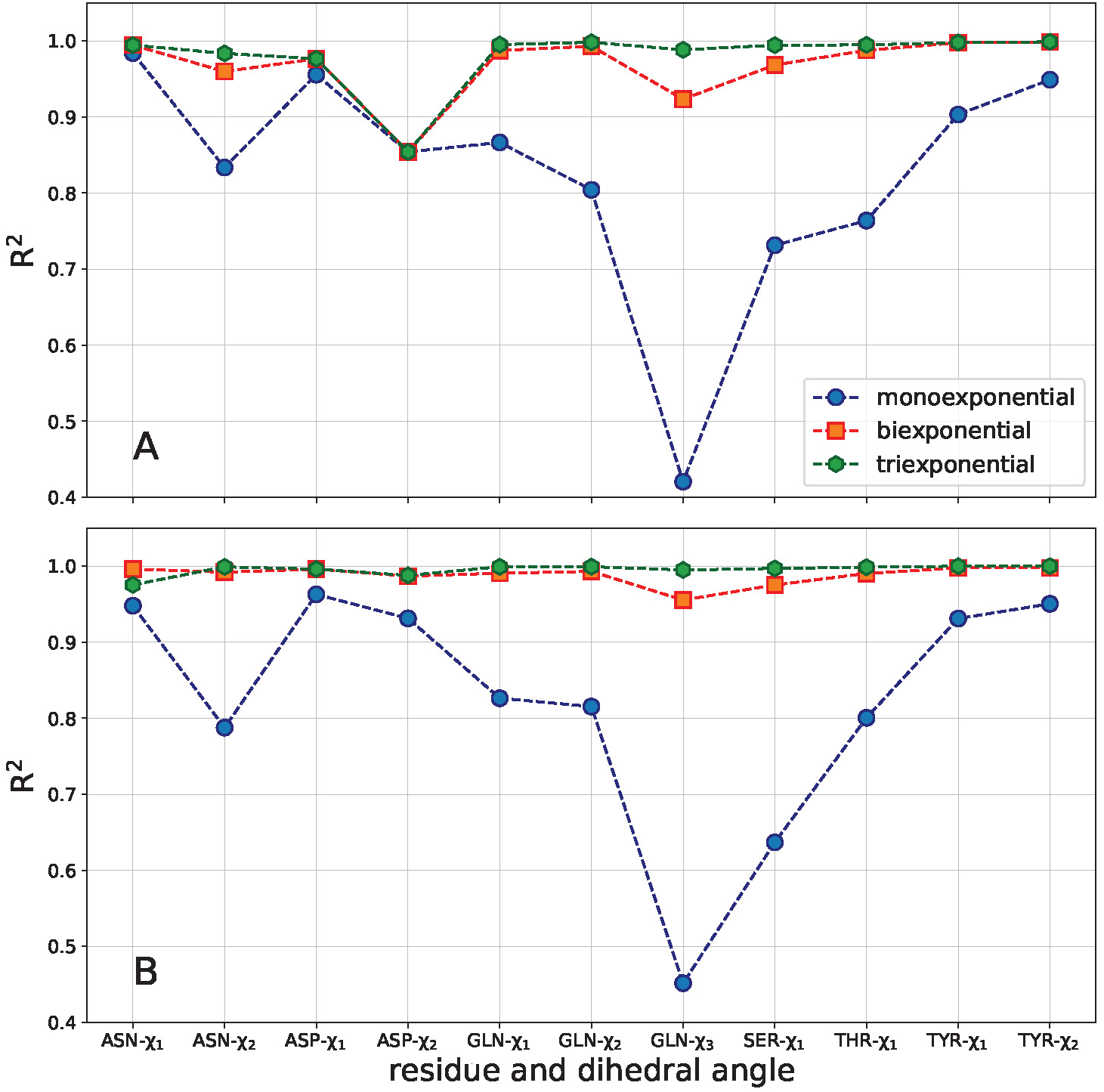
Coefficient of determination (R^2^) for mono-, bi-, and triexponential fits of the ACFs of dihedral rotations in the (A) dilute and the (B) condensate systems.

